# Reliability-Based Voxel Selection

**DOI:** 10.1101/703603

**Authors:** Leyla Tarhan, Talia Konkle

**Affiliations:** Department of Psychology, Harvard University, 33 Kirkland St., Cambridge, MA 02140, United States of America; Center for Brain Science, Harvard University, 33 Kirkland St., Cambridge, MA 02140, United States of America

**Keywords:** fMRI, voxel selection, data reliability, condition-rich designs

## Abstract

While functional magnetic resonance imaging (fMRI) studies typically measure responses across the whole brain, not all regions are likely to be informative for a given study. Which voxels should be considered? Here we propose a method for voxel selection based on the reliability of the data. This method isolates voxels that respond consistently across imaging runs while maximizing the reliability of multi-voxel patterns across the selected voxels. We estimate that it is suitable for designs with at least 15 conditions. In two example datasets, we found that this proposed method defines a smaller set of voxels than another common method, activity-based voxel selection. Broadly, this method eliminates the need to define regions or statistical thresholds *a priori* and puts the focus on data reliability as the first step in analyzing fMRI data.

## 1. INTRODUCTION

As functional magnetic resonance imaging reaches into its third decade of use in cognitive neuroscience, researchers continue to develop new ways to analyze patterns of activity in the brain. Current tools range from univariate contrasts to Representational Similarity Analysis (Kriegeskorte, 2008), decoding (Cox & Savoy, 2003; Hanson et al., 2004; Norman et al., 2006), encoding models (Mitchell et al., 2008; Huth et al., 2012), clustering (Lashkari et al., 2010), and more. These analyses vary in complexity and the types of questions they answer. But in all cases, the researcher must first answer the same question: where in the brain should these analyses be done?

The possible answers to this question fall along a spectrum between focusing on specific regions of interest (ROIs) and considering every voxel in the brain. ROI-based research (e.g., Johansen-Berg et al., 2004; Saxe et al., 2006) typically restricts all analyses to an area of cortex defined using separate functional or anatomical localizers. ROI approaches make two assumptions: that these voxels are interesting regions of cortex to investigate, and that they might make up a homogenous unit, serving the same cognitive functions (though these assumptions can be tested; e.g., see Duncan & Owen, 2000; Jiang & Kanwisher, 2003). As a benefit, targeted ROI analyses circumvent the statistical challenge of correcting for multiple comparisons, which occurs when analyses are done at the whole-brain level.

In contrast to an ROI-based approach, whole-brain approaches (e.g. whole-brain contrasts, information mapping techniques, and searchlight analyses; Norman et al., 2006; Kriegeskorte et al., 2006) have the advantage of not relying on pre-conceived ideas about where in the brain one might expect an effect. However, these analyses require stringent corrections for multiple comparisons, which can obscure all but the strongest effects, running counter to the goal of obtaining and analyzing data at a fine spatial scale. Additionally, searchlight analyses, which are effectively roving ROI analyses using spheres of voxels (Norman et al., 2006; Kriegeskorte et al., 2006), also require implicit pre-conceived hypotheses about neural coding: researchers must have a hypothesis about the spatial scale at which an effect will be observed, and also make the assumption that these mini-regions-of-interest are spherical.

One additional challenge for whole-brain approaches is that they do not address the issue of variable signal quality over the brain. It is well known that the signal measured in fMRI varies significantly over the cortex, and in some cases carries a troubling amount of noise (Eklund et al., 2016). Further, the quality of the signal may depend upon the task – for example, areas in the fronto-parietal attention network might respond more consistently during an attentionally-demanding task than when freely viewing an image. Therefore, another approach that sits between the ROI-based and whole-brain extremes is to isolate a relatively broad swathe of cortex with a high-quality signal, such as visually-responsive voxels within occipito-temporal cortex.

Unlike more classic regions of interest, a broad swathe approach does not assume that these selected voxels make up a singular functional unit *per se*; rather, it assumes that responses in these voxels are more relevant to the researcher’s analysis plan and hypothesis than the rest of the brain. Within these broad regions, analyses can be done to characterize the large-scale structure of response preferences (e.g., Hasson et al., 2002; Hasson et al., 2003; Orlov et al., 2010; Konkle & Oliva, 2012), or to evaluate encoding models (e.g. Mitchell et al., 2008; Tarhan & Konkle, 2019), or to assess population coding (Haxby et al., 2011; Haxby, 2012). The benefit of this approach is that the cortex under consideration is more extensive than a small ROI, while still being relevant to the theoretical questions at stake. However, a challenge for this approach is that it is not obvious how to select these high-quality voxels in the first place. Different researchers have approached this challenge in different ways.

First, one popular method is to select the voxels that are most “active” in response to the stimuli (Kriegeskorte et al., 2008; Konkle & Oliva, 2012; Mur et al., 2013; Jozwick et al., 2016; Kay et al., 2017). This responsiveness is often measured with a contrast between all stimulus conditions and rest: voxels with a *t*-value greater than some pre-set bound (e.g., *t*>2) are considered “active.” In other words, this method isolates regions that respond positively on average across stimulus conditions, relative to baseline. Activity-based voxel selection is sensitive to the signal strength of a given voxel because it is based on *t*-values, which are low if noise is high; however, this method is *not* sensitive to systematic differences across conditions, as voxels that respond equally across all conditions could still be considered active. Given that this is a popular choice in many recent fMRI papers – especially those studying the visual cortex, as we do – we treat the activity-based voxel selection method as the main comparison for the reliability-based method described here.

Another voxel-selection approach identifies voxels that maximize the variance among conditions (Pereira et al., 2009), or that are well-fit by a candidate feature model (Nishimoto et al., 2011; Naselaris et al., 2012, Guclu & van Gerven, 2015; Huth et al., 2016). This approach makes sense within a modeling framework, but in most cases requires a pre-specified threshold for “well-fit” voxels. Relatedly, other researchers have focused on selecting voxels that respond stably across conditions. For example, Mitchell et al. (2008) used a cross-validation procedure to isolate voxels that respond similarly across folds of the data. Similarly, Norman-Haignere et al. (2015) selected voxels that were both active (showed a significant response to sounds vs. silence) and responded consistently across scans. These selection methods are sensitive to the activation profile of each voxel, but are also limited because they often rely on a pre-set statistical threshold.

The current paper introduces a procedure for voxel selection that starts from a similar logic as this last method, but instead selects voxels based on their *reliability* across scans. Specifically, in the first step, the reliability of every voxel is computed based on the voxel’s profile of responses to all conditions in two halves of the data. Next, a range of voxel-reliability thresholds are considered, and the stability of the multi-voxel patterns within the voxels that survive each threshold is assessed for each condition. Finally, the researcher can select a reliability cutoff to select the final set of voxels, while explicitly being able to balance the tradeoffs between maximizing spatial coverage on one hand and multi-voxel pattern reliability on the other. This procedure forms the first step in an fMRI analysis: once the reliable voxels are selected, all subsequent analyses can be conducted in only these voxels.

Our method differs from previous stability-based approaches in two key ways. First, the process of identifying an acceptable reliability threshold is done based on the data, without specifying a parameter value *a priori*. Second, our method selects voxels whose responses vary across conditions; that is, voxels that respond more to some conditions than to others. This contrasts with the method used in Norman-Haignere et al. (2015), which includes regions that respond similarly to all conditions. Thus, our method is especially well-suited to analyses that leverage the variance in the data, for example to predict brain responses or representational dissimilarities using encoding models.

Here, we illustrate the use of this reliability-based voxel selection procedure on a typical set of condition-rich fMRI data: whole-brain responses to 60 everyday actions (Tarhan & Konkle, 2019). While this method was originally developed for isolating voxels for a subsequent voxel-wise encoding analysis, we also demonstrate how it might be used with other analyses, such as classification-based approaches. Further, we replicate all analyses in a separate dataset of responses to 72 everyday objects (Magri et al., 2019). Finally, we discuss the benefits and limitations of this method, and possible use cases going forward.

## 2. MATERIALS AND METHODS

### 2.1 Dataset Description

The primary dataset used in this paper consists of whole-brain functional responses to 60 everyday action videos, collected from 13 human subjects (Figure 1a). Participants completed four functional runs, during which they freely viewed the 2.5-second videos and detected an occasional red frame to maintain alertness. Functional and anatomical data were pre-processed using Brain Voyager QX software (Brain Innovation, Maastricht, Netherlands). General linear models (GLMs) were fit to data from odd and even functional runs with a regressor for each condition (one video). Each voxel’s timecourse was first z-transformed within an fMRI run, then corrected for temporal autocorrelations. The beta weights were extracted from whole-brain GLMs, yielding estimates of each voxel’s response to each condition. This was done separately for each participant, in addition to a whole-brain random effects GLM to quantify responses at the group-average level. More information can be found in Tarhan & Konkle (2019). For replication purposes, we also included a supplementary dataset collected from 11 human subjects viewing still images of 72 everyday objects (Magri et al., 2019). These data were pre-processed following similar procedures. Both datasets can be downloaded from the Open Science Framework (https://osf.io/m9ykh/). In order to perform reliability-based voxel selection, the only necessary components of these data are odd- and even-run beta values for each condition in every voxel.

**Figure 1:**
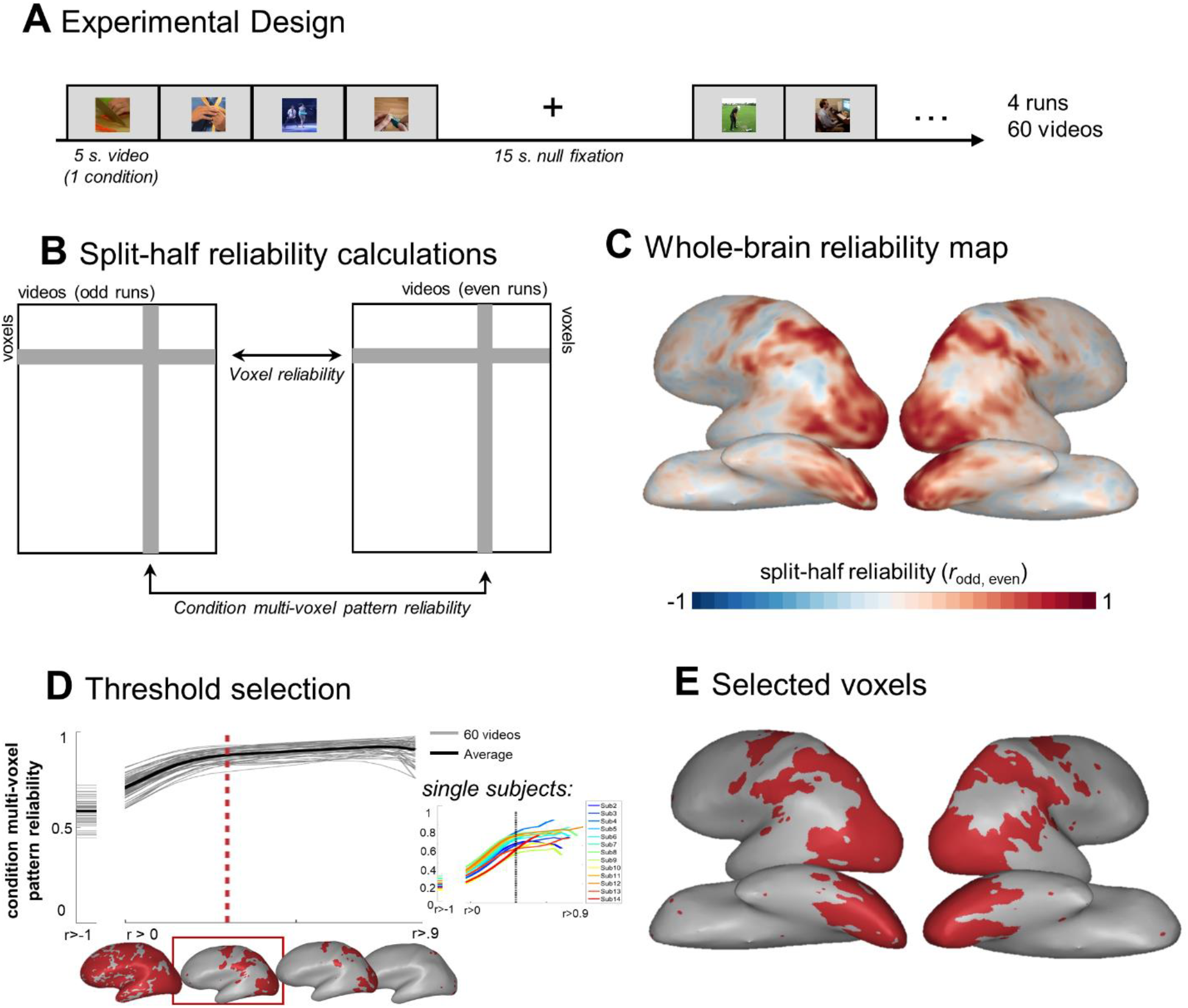
Summary of the Reliability-Based Voxel Selection Procedure. (a) Schematic of the experimental design for the main dataset: participants viewed 60 everyday action videos while detecting an occasional red frame to maintain attention. (b) Split-half reliability was computed for all voxels by correlating their beta weights for all 60 conditions across even and odd runs. Condition multi-voxel pattern reliability was computed for all conditions (videos) by correlating beta weights over all voxels across even and odd runs. (c) Whole-brain map of split-half voxel reliability in this dataset. (d) Curve showing the pattern reliability for each condition (y-axis) at a range of possible voxel inclusion thresholds (x-axis). Red line indicates the final threshold selected for this dataset (*r* = 0.30). Inset figure shows the same graph as in (d) but computed for individual subjects. Each line is the reliability curve for the average pattern reliability across conditions for one participant. (e) Voxels that survived the final inclusion threshold are shown in red.

### 2.2 Reliability-Based Voxel Selection

Reliability-based voxel selection proceeded in two steps, summarized in Figure 1. First, split-half reliability was calculated separately for each voxel in the brain by correlating each voxel’s *response profile* (the vector of beta weights in response to each condition) across even and odd functional runs (Figure 1b). This produced a whole-brain reliability map, with one *r-*value for each voxel in the cortex (Figure 1c). Second, to determine which voxels to include in subsequent analyses, we considered a continuum of possible thresholds (from *r*>0 to *r*>0.95). At each threshold, we calculated the reliability of each condition’s *multi-voxel pattern* (i.e. the pattern of responses to a single condition) in voxels that survived that threshold. *Condition multi-voxel pattern reliability* was calculated for each condition by correlating its multi-voxel response pattern across even and odd functional runs. This was done separately for each condition, and then averaged across conditions. This analysis produced a smooth curve, where the average condition multi-voxel pattern reliability increases as more reliable voxels are included in the selection (Figure 1d).

This curve also illustrates a natural tradeoff between the coverage and the stability of the data – simply choosing the threshold that produces the highest possible condition-pattern reliability limits coverage to a tiny fraction of the cortex (Figure 1d). While the data in that fraction will be highly reliable, this strategy runs counter to experimental goals aimed at studying the large-scale structure of the cortex. On the other hand, a threshold that is too lax may include regions with erratic responses. Therefore, an optimal threshold for voxel inclusion balances coverage and the stability of the data, but its exact value depends on the goals of the experimenter. This curve serves as a guide to balance these tradeoffs. In our case, in multiple datasets we observed that this curve has a plateau – after some critical point, increasing the inclusion threshold only minimally increases the data’s reliability but continues to limit coverage. Therefore, we took this plateau point to be an appropriate inclusion threshold.

How should one select this point? To automatically detect its location, one could objectively locate the point where the curve’s second derivative equals zero, which intuitively is when the slope begins to level off. However, we recommend following a more heuristic method. We selected the plateau point by eye, and then considered a range of nearby thresholds to select one that provides good coverage of our general regions of interest while minimizing small, extraneous groups of voxels (“speckles”).

Critically, notice that this threshold decision is made independently of any hypotheses about how a given region or voxel will respond to the conditions. Instead, this voxel selection procedure only depends on the fact that voxels respond consistently across scans but vary in how much they respond across conditions; it depends not at all on *which* conditions it responds to the most. Because of this, reliability-based voxel selection is independent of any particular hypotheses regarding the relationship among conditions.

Another benefit of this method is that it is fairly flexible; for example, it can be applied at the level of the group (Figure 1d) or individual subjects (Figure 1d, inset). Interestingly, in both datasets we found that the reliability plateaued at a very similar point across participants. Therefore, we selected a reliability cutoff based primarily on the group-level data, then applied that cutoff to the single-subject data to define reliable voxels separately in each subject. While this cutoff was similar across subjects, the number of voxels that survived it varied across subjects. We explore the consequences of this variability further in the **Discussion**.

Code implementing these procedures can be downloaded from the Open Science Framework (https://osf.io/m9ykh/).

## 3. RESULTS

### 3.1 Reliability based voxel selection

We performed reliability-based voxel selection on our primary sample fMRI dataset, in which participants viewed short videos of people performing a wide range of actions. Based on the reliability curve shown in Figure 1d, we selected an inclusion threshold of *r* = 0.30. Figure 1e displays the voxels that survive this threshold. The selected voxels reveal extensive coverage of both the ventral and dorsal visual streams as well as primary somatosensory and motor cortices, while excluding less-reliable regions in the anterior and superior temporal lobe and the pre-frontal cortex.

### 3.2 Comparing Reliability- and Activity-Based Voxel Selection

For comparison with this reliability-based selection method, we also considered a voxel-selection method based on overall activity. First, a *t*-test was conducted over the contrast of all 60 conditions > rest in every voxel. Second, voxels with a *t*-value greater than 2.0 were selected (Figure 2a, blue voxels). Researchers employ diverse *t*-thresholds to define active voxels. While some choose an arbitrary cutoff such as 2.0 (Long et al., 2018) or 0 (Kay et al., 2017), others select the *n* voxels with the highest *t*-value, where *n* is a somewhat arbitrary size such as 100 (Kriegeskorte et al., 2008; Mur et al., 2013; Jozwick et al., 2016). Based on a student’s *t-*distribution, *t*=2.0 roughly corresponds to a significance level of *p* < 0.05 without corrections for multiple comparisons, making it a relatively liberal threshold.

**Figure 2:**
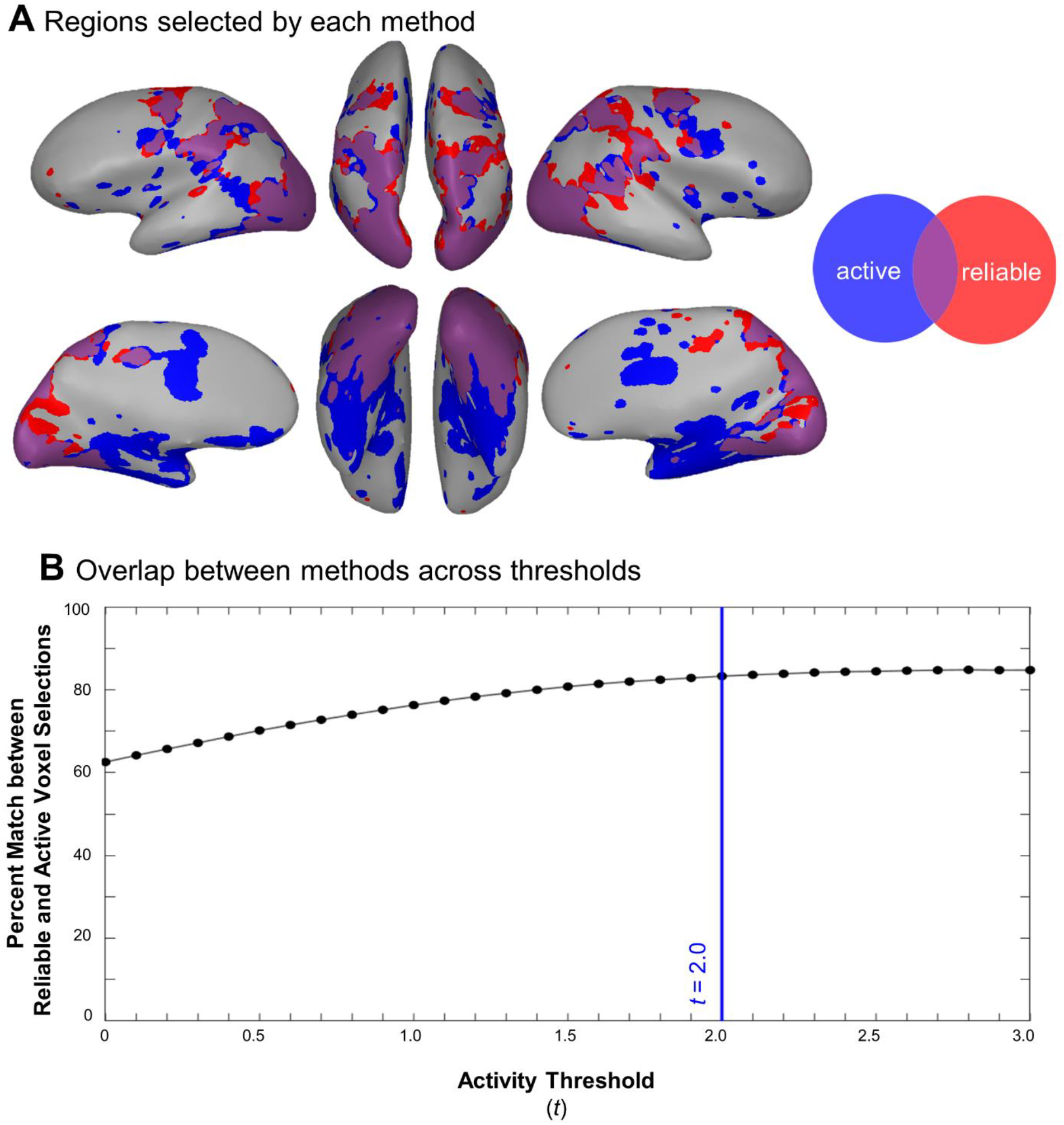
Comparison between the Data Selected Using Reliability and Activity. (a) Selected voxels from both methods are plotted on an example subject’s inflated cortex. Red voxels are reliable, blue voxels are active, and purple voxels are reliable and active. Voxel selection is shown for the group data. (b) Results of the analysis comparing reliability- and activity-based voxel selections at a range of possible *t*-thresholds. At each threshold, the percentage of voxels in the gray matter that were categorized in the same way (selected or not selected) by both voxel selection methods is plotted. Blue line indicates the threshold used to define active voxels in the main analyses (*t* = 2.0).

Do these methods result in coverage of different brain regions? Figure 2a shows both voxel selections, including active voxels (blue), reliable voxels (red), and their overlap (purple). In general, the reliability-based selection method isolated a smaller set of voxels that are more localized to the occipito-temporal cortex (*t*(12) = 7.12, *p* < 0.001).

Given that the choice to use an activity threshold of *t >* 2.0 was arbitrary, it is natural to ask whether these results change if active voxels are defined using a different threshold. In other words, is there an activity threshold such that active and reliable voxels cover the same regions? After considering a range of thresholds between *t >* 0 and *t >* 3.0, we found that the degree of overlap between active and reliable voxels followed a curve that plateaued around 80% at *t* > 2.0 (Figure 2b; **Supplemental Methods**). Thus, in this dataset, choosing a different activity threshold would not dramatically increase the degree of overlap between reliable and active voxels beyond what we observe here. In summary, when using standard thresholds, the activity-based selection method included more voxels than the reliability-based method.

### 3.3 How many conditions are necessary?

Reliability-based voxel selection is well-suited to condition-rich fMRI designs, which expose subjects to many conditions (or items). With very few conditions, the estimates for a voxel’s split-half reliability will not be stable, as a correlation based on only a few points is not a robust measurement of a linear relationship (Bonnett & Wright, 2000). How many conditions must one test to employ this voxel selection method?

To answer this question, we performed a simulation analysis based on our example data. We asked how robust the split-half reliability calculation is at a range of numbers of conditions (from 1 to 60). For each possible number of conditions *c*, we randomly selected responses to *c* conditions, then calculated each voxel’s split-half reliability. After doing so 100 times, we calculated the standard deviation of each voxel’s split-half distribution. Figure 3 shows the average standard deviation across voxels for each number of conditions *c.* In general, stability improves (standard deviation falls) as the number of conditions grows. However, the pattern is not linear: at a certain point, adding more conditions does not confer additional benefits to stability. In our data, this point occurred at 15 conditions. This suggests that the reliability-based voxel selection method is appropriate for designs measuring responses to 15 or more conditions.

**Figure 3:**
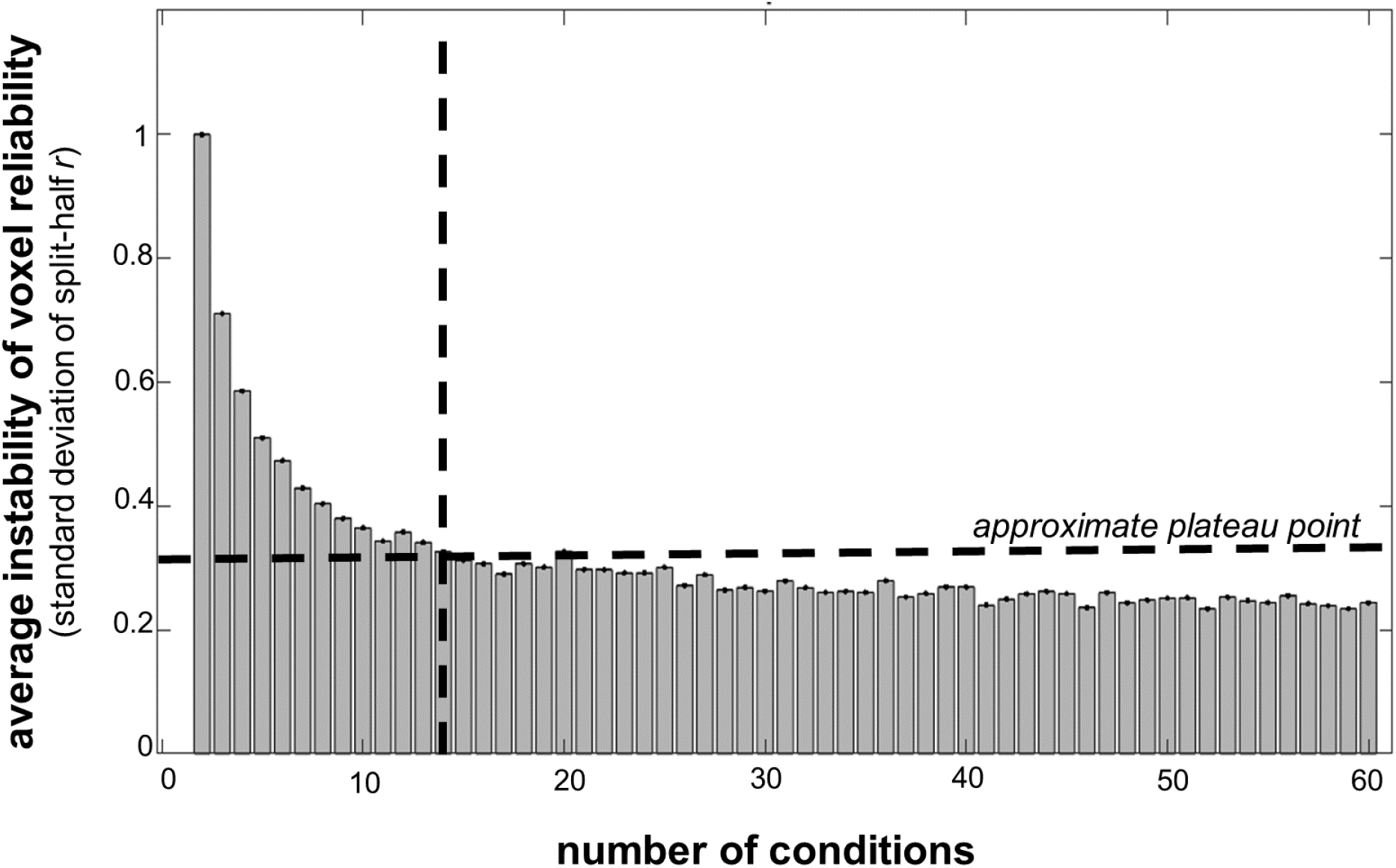
Estimating the Minimum Number of Conditions for Reliability-Based Voxel Selection. Average instability of the voxel reliability calculation is plotted for every number of conditions *c* from 1 to 60. For each *c*, we selected a random subset of whole-brain responses to *c* conditions. We then calculated the split-half reliability for every voxel within those subsetted data. This procedure was repeated over 100 iterations. Instability was calculated for each voxel as the standard deviation of split-half reliability over the 100 iterations, then averaged across voxels. Crosshairs indicate the approximate point at which the tradeoff between number of conditions and instability begins to plateau (15 conditions), meaning that this is the minimum number of conditions needed to use this reliability-based voxel selection method.

However, there is an important caveat to this result. In this data set the stimuli are sampled broadly from the action domain, with extensive variability across stimuli. We have not tested this method on other kinds of condition-rich designs using stimuli that may be more homogenous (e.g., only faces) or that have a more categorical distribution (e.g., 10 faces and 10 places). Thus, the number of conditions indicated here should not be treated as a strict recommendation; instead, we encourage researchers to test this in their own pilot data.

### 3.4 Replication

These findings replicated in an independent dataset of responses to 72 real-world objects (Figure 4a). Figure 4b shows the split-half reliability for this dataset over the brain. Based on the reliability curve shown in Figure 4b, we selected a reliability cutoff of *r* > 0.30. Although it is striking that we found the same cutoff in both datasets, we assume that this is mere coincidence. As we found in the first dataset, the reliability-based method selected fewer voxels than the activity-based method (*t*(10) = 3.89, *p* < 0.01; Figure 4c). In the temporal, parietal, and lateral occipital cortices, reliable voxels were primarily a subset of active voxels. Whereas for the most part reliable voxels were a subset of active voxels in the action video dataset, in this dataset early visual regions in the medial occipital cortex contained extensive reliable voxels but very few active voxels. Finally, split-half reliability estimates began to stabilize around 15 conditions, further corroborating our estimate that it is necessary to include at least 15 conditions to employ reliability-based voxel selection, given a broadly sampled domain.

**Figure 4:**
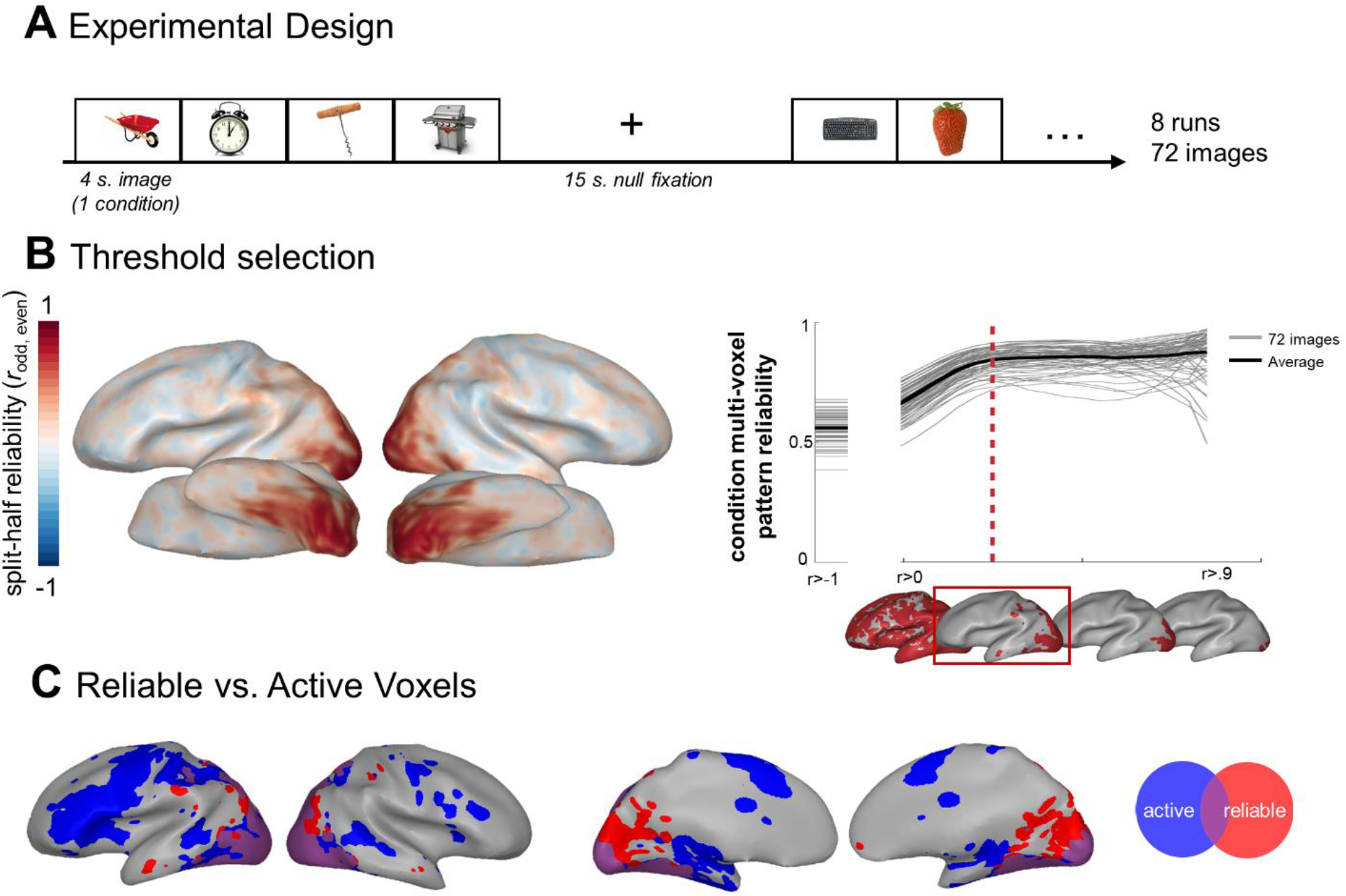
Replication. (a) Schematic of the experimental design for the replication dataset: participants viewed images of 72 everyday objects while detecting an occasional red frame to maintain attention. (b) Whole-brain map of split-half voxel reliability in this dataset and curve showing the pattern reliability for each condition (y-axis) at a range of possible voxel inclusion thresholds (x-axis). Red line indicates the final threshold selected for this dataset (*r* = 0.30). (c) Comparison of the voxels selected based on reliability and activity. Selected voxels from both methods are plotted on an example subject’s inflated cortex. Red voxels are reliable, blue voxels are active, and purple voxels are reliable and active. Voxel selection is shown for the group data.

### 3.4 Decoding Analysis

As noted above, this voxel selection method was initially designed to be used in voxel-wise encoding modeling analyses, where the aim is to map out the functional properties of the cortex by measuring individual voxels’ tuning to particular features (e.g., Tarhan & Konkle, 2019). However, given the importance of analyzing reliable brain data, it stands to reason that reliability-based voxel selection may also be a useful tool for analyses that aggregate over multiple voxels to characterize a whole region, such as decoding or representational similarity analysis. What steps are necessary to adapt this method to these kinds of analyses? And does using more reliable voxels lead to better outcomes? To explore these questions, we performed pairwise classification to decode action identity from reliable voxels. We then assessed the decoding accuracy at a range of reliability thresholds and asked whether accuracy was higher among more reliable voxels (see **Supplemental Methods**).

This decoding analysis required an adjustment to the reliability-based voxel selection procedure: for analyses that aggregate over multiple voxels, it is important to use *different* data to define the reliable voxels and perform the decoding analysis. Otherwise, the results may be biased by the existence of “lucky” noise, which is correlated across odd and even runs. Therefore, for each pair of conditions that was decoded, we selected the reliable voxels using responses to all conditions *except* that pair. Then, the decoding was done within those reliable voxels (**Supplemental Methods**; Figure S2a & b). Note that one consequence of this procedure is that each pair of conditions was decoded based on responses from a slightly different set of voxels.

Figure 5 shows the average decoding accuracy as a function of the voxel-reliability threshold. Accuracy increased when restricting the analysis to voxels with higher reliability up to a threshold of *r* = 0.60 (accuracy = 87.2%), and interestingly then began to fall around *r* = 0.7. Note that this drop in accuracy is not due to a catastrophic loss of coverage – over 1,000 voxels survived the threshold at *r* = 0.7 (see coverage maps below the x-axis). For comparison, the red dashed line indicates our reliability-based voxel-selection threshold, which is at or near the point where this accuracy curve plateaus. This pattern replicated in the second dataset (responses to 72 real-world objects; Supplemental Figure 2). Thus, reliability-based voxel selection can yield reasonable decoding performance, even though the outcome measure is not directly leveraged during voxel selection. However, selecting reliable voxels does not guarantee coverage in regions that represent the feature being decoded, so simply maximizing reliability does not necessarily maximize decoding performance.

**Figure 5:**
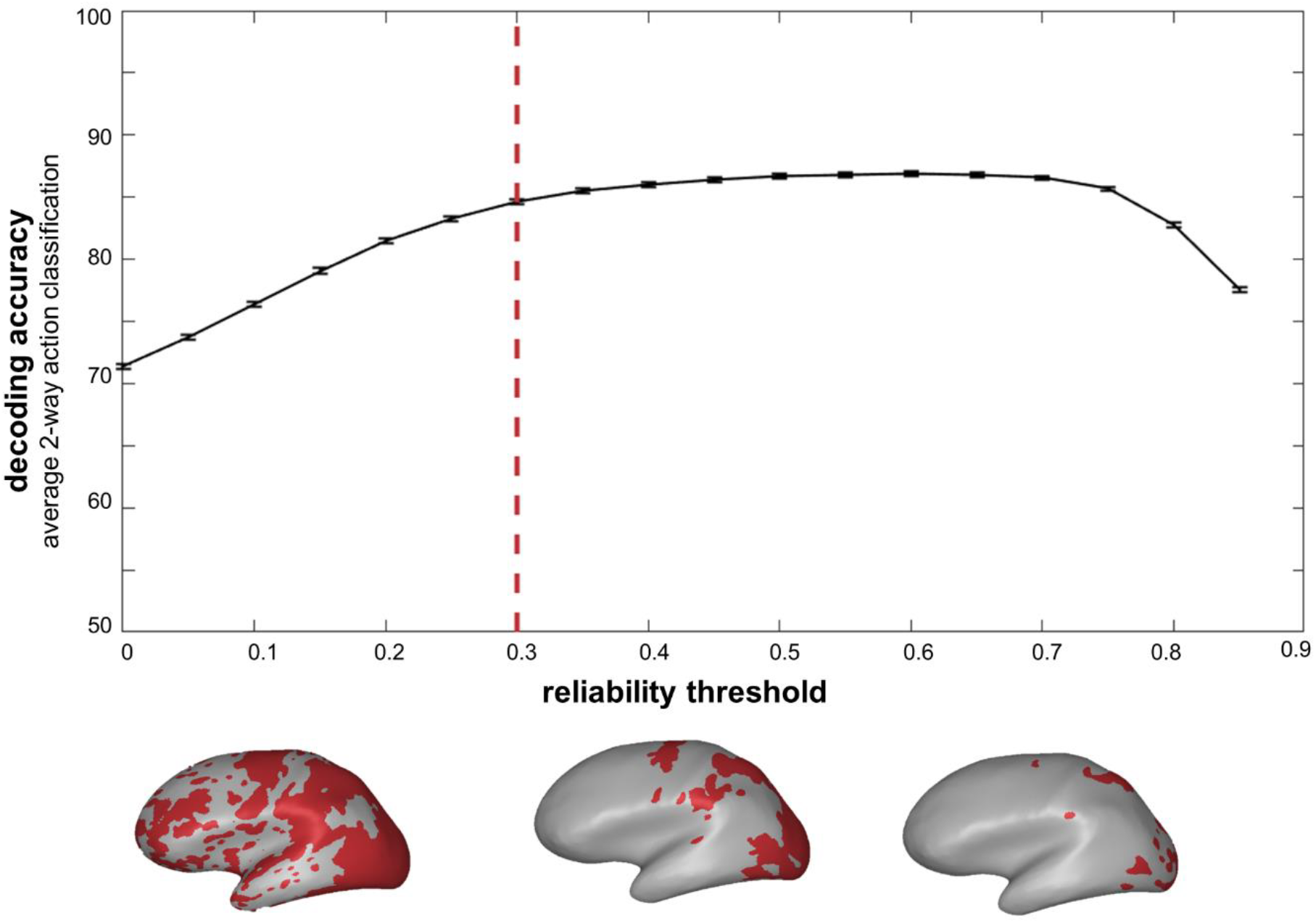
Decoding Accuracy. Results of the pairwise action decoding analysis are plotted at a range of reliability thresholds. The black curve indicates average performance across action pairs, and error bars indicate the standard error of the mean. Brain figures show the reliable coverage for the group data at three sample thresholds: *r* = 0.1, 0.4, and 0.7. Red dashed line indicates the final reliability threshold selected for this dataset (*r* = 0.3).

## 4. DISCUSSION

The method we have outlined here introduces a new way to select voxels from the whole brain that respond systematically to experimental variations. In addition, it is straightforward to implement and is informed by the data, rather than by *a priori* parameters that specify a statistical cut-off or a fixed region size. We found that this method selects voxels with more limited coverage than activity-based selection. Once selected, reliable voxels could be entered into a wide range of hypothesis-driven analyses; therefore, we suggest that reliability-based voxel selection is a promising first step to make subsequent fMRI analyses more robust.

### 4.1 What properties differentiate reliable and active brain regions?

Given that activity- and reliability-based voxel selection result in different sets of voxels, are there interesting differences in the neural signals present in these regions? One possibility is that, while activity-based voxel selection takes noise into account to some extent, some active voxels are still more noisy than reliable voxels. Another possibility is that “active” regions are less sensitive to the condition structure in the data. Whereas reliability requires some variance among conditions, a region can be “active” even if it responds with equal magnitude to all conditions. Therefore, if a researcher predicts that equal response magnitudes across all conditions would be meaningful – for example, if she hypothesizes that a scene-processing region will respond equally across all outdoor and indoor scenes – overall activity might be a reasonable way to select voxels. In contrast, a researcher using an analysis that leverages the variance in the data, such as encoding modeling, representational similarity analysis, or most univariate analyses, should select regions whose responses vary across conditions.

### 4.2 Comparison with Outcome-Based Methods

Apart from activity-based selection, one of the main alternatives to reliability-based voxel selection is to identify voxels based on the outcome of an analysis; for example, to identify voxels where a model is well-fit. While such outcome-based methods are a useful tool in some situations, reliability-based selection offers two major advantages that make it more generally applicable. First, reliability-based selection may identify regions that have sufficient variance to be included in the analysis, but nevertheless do not show the hypothesized effect. This is informative because it allows researchers to conclude something about the representations in both the regions where the effect exists (they match the prediction), and those where it doesn’t (they have some other format, which is still reliable). In contrast, with an outcome-based voxel selection method it would be impossible to conclude anything about the unselected regions: they could show no effect, or they could simply contain bad data. Second, reliability-based voxel selection makes it easy to select regions for multiple kinds of analyses. For example, if a researcher plans to use model comparisons to determine which candidate features fit the data the best, which model should she use to select her voxels? Reliability-based voxel selection avoids this situation by divorcing the selection procedure from the analysis outcome.

### 4.3 Double-dipping and Bias

Double-dipping occurs when a voxel selection procedure biases subsequent analyses in the direction of the hypothesis by capitalizing on favorable noise (e.g. Kriegeskorte et al., 2009). For example, if an ROI is defined based on the contrast viewing pictures of faces > viewing pictures of objects, then the estimate of how much more this ROI responds to faces than objects will be biased if assessed with the same data that were used to define the ROI. In such a case, it is important to use separate datasets to define and test the ROI, to avoid inflating or even fabricating effects due to noise in the direction of the hypothesis. In contrast, our method merely finds voxels that reliably respond more to some conditions than others, without any reference to *which* conditions those are (e.g., Norman-Haignere et al., 2015; Mitchell et al., 2008). Thus, any subsequent assessment that *maps out* the functional properties of the cortex, such as mapping peak response conditions or predicting voxel-responses in an encoding-modeling framework, is unbiased by the voxel selection.

To illustrate this more intuitively, suppose participants saw images of many different objects. As experimenters, we may be interested in whether any voxels respond parametrically based on the real-world size of the objects, or how graspable the objects are, or how aesthetically pleasing the objects are. We could reasonably use the whole dataset to define a set of reliable voxels in which to run such an analysis, because there is no guarantee that the selected voxels will respond according to our dimensions of interest. Some might respond more to big than small objects, and others might respond more to redder than bluer objects. The selection procedure does not bias an analysis searching for these hypothesized representations. Thus, researchers can leverage the entire data set and use reliability-based voxel selection as a first step after collecting condition-rich fMRI data, as this step is completely separate from any subsequent hypothesis-driven analyses.

In contrast to mapping analyses, some analyses aim to read information out from multi-voxel patterns (e.g., classification or RSA). For these read-out analyses, it is no longer safe to use the same data to define reliable voxels and analyze their responses. The reason for this is that some voxels could be reliable in both halves of the data by chance alone. Including these voxels introduces “lucky” noise that is shared across halves of the data, which a classifier may capitalize on. As Figure S2c illustrates, this mechanism boosted decoding accuracy when we used the full dataset to define the reliable voxels, rather than held-out data (particularly at low reliability thresholds).

To avoid this bias, it is necessary to use different data to define reliable voxels and analyze the data when analyzing multi-voxel response patterns. This can be done in several ways. One option is to use completely separate datasets to define reliable voxels and analyze their responses; though, splitting a dataset so many ways can cause power issues. A second option is to use the full dataset to determine the best reliability-threshold to define reliable voxels, then iteratively select reliable voxels during the subsequent analysis while holding out the pair of conditions being decoded (as we did in the decoding analysis discussed in the **Results**).

Finally, it is also important to note that, while selecting voxels based on reliability does not bias subsequent hypothesis-driven analyses based on differences between stimuli, the selection procedure could bias estimates of an effect’s variance (i.e., errorbars or significance) if those estimates incorporate the variance of responses across repetitions. Thus, all estimates of variance and significance should be performed across stimuli or subjects, rather than across *repetitions* of the same stimuli.

In sum, reliability-based voxel selection can be used with a large variety of analyses—however, it is critical to consider the eventual aim of the analysis it is paired with before deciding on its exact implementation. If the aim is to map the properties of individual voxels, the same data can be used to define reliable voxels and conduct the mapping analysis. However, if the aim is to characterize the multi-voxel patterns in a larger region, it may be necessary to use cross-validation or a separate dataset to select the voxels and analyze the data.

### 4.4 Novel Applications

This voxel selection method also allows for several variants, which highlight the many facets of reliability. Data can be reliable over different sources of variation – while we calculated split-half reliability between odd and even fMRI runs (variation in time), one could split the data in other ways to isolate meaningfully-different brain regions. For example, if an experiment includes two stimulus sets – perhaps set 1 contains photographs of real-world objects and set 2 contains drawings of the same objects – one could use this method to find regions that respond reliably across the two sets. This would isolate regions that are robust to low-level variations in stimuli, suggesting that their responses reflect higher-level properties of the stimuli, such as object shape or identity. One could also calculate reliability across subject groups – for example, to identify regions that respond consistently to hearing the names of object in blind and sighted participants. This variant of the method would imply that the responses of the selected regions in the sighted may not solely be related to visual responsiveness. In a different vein, one could improve the reliability of region-of-interest analyses by selecting the reliable voxels that lie within a region of interest or functional parcel (Federenko et al., 2010; Julien et al., 2012). It is also possible to calculate the reliability of distances between conditions, within a representational similarity framework (Thornton & Mitchell, 2017). In general, requiring reliable responses over different manipulations—be they at the stimulus, task, or subject group level—can help to isolate the most relevant brain regions for the theoretical question at hand.

### 4.5 Selecting Voxels at the Single-Subject and Group Levels

In our data, we found voxel inclusion thresholds that were highly consistent across subjects and the group data. However, this does not guarantee that reliable voxels surviving those thresholds will overlap perfectly across subjects. In the primary dataset examined here, subjects’ reliable coverage using the same cut-off varied from 1,596 to 6,879 voxels with a 3×3×3 mm resolution. Thus, the choice to use this voxel selection method at the level of single subjects or the group depends upon the kind of question being asked.

For example, in a single-subject approach, one could define reliable voxels separately in each subject, and then perform single-subject analyses in those voxels. This approach is particularly useful if the question concerns the link between an individual’s experience, traits, or judgments and their neural responses. However, this approach may force the experimenter to analyze different brain regions in different subjects. An alternative approach is to do a group-level analysis, defining reliable voxels based on the group data, then only analyze the group data.

In many cases, a hybrid analysis scheme may be appropriate: the voxels are defined using the group data, and all single-subject analyses can be performed within that set of voxels. This approach guarantees that the same regions will be examined across subjects, enabling the researcher to compare response maps between subjects.

However, some of the data will inevitably come from less reliable voxels. Therefore, it is important to interpret each subject’s results in light of their reliable coverage.

## 5. CONCLUSIONS

In summary, reliability-based voxel selection is a principled method for isolating regions of the brain for further analysis that is informed by the stable structure in the dataset. The strengths of this method are that (i) it is straightforward to implement, making it easy to adopt into a variety of fMRI analysis settings, (ii) it is agnostic to *a priori* hypotheses about which conditions drive which cortex where, and (iii) it puts the emphasis on data reliability as an early step in fMRI data analysis processing.

## ACKNOWLEDGEMENTS

We are grateful to several extremely thoughtful and careful reviewers for improving the accuracy and scope of this work. Thank you also to Emilie Josephs and the Cambridge Writing Group for their support during the writing process. Funding for this project was provided by NIH grant S10OD020039 to Harvard University Center for Brain Science, NSF grant DGE1144152 to L.T., and the Star Family Challenge Grant to T.K.

## Supplemental Information

### Extended Methods

#### Activity-based Voxel Selections with Different Thresholds

In the main analysis, the activity-based threshold was set at *t* > 2.0. Here, we considered a range of additional thresholds between *t* > 0 and *t* > 3.0. At each threshold, we evaluated the match between the reliable and active voxel selections by calculating the number of voxels that were categorized in the same way (either selected by both methods or ignored by both methods). We then calculated the percent of voxels in the gray matter that met this criterion (Figure 2b).

#### Decoding Analysis

To determine the effect of reliability-based voxel selection on the outcome of an analysis, we performed pairwise classification to decode action and object identity at a range of reliability thresholds. We considered every threshold from *r* > 0 to 0.95 at intervals of 0.05, but only analyzed data for thresholds that were survived by at least 10 voxels. Reliable voxels were defined separately for each pair of conditions, using the group average responses to all conditions except the pair being classified. At each threshold, we iteratively trained a linear support vector machine classifier to distinguish between all condition pairs on 80% of the data, then tested the classifier on the remaining 20% of the data by calculating the percentage of the test responses that were correctly classified. The data for each condition consisted of odd- and even-run responses from each subject. This was done across 100 iterations. Average classification accuracy (averaged across both condition pairs and iterations) is plotted at every threshold in Figure 5 for the primary dataset and Figure S1 for the replication dataset.

**Figure S1:**
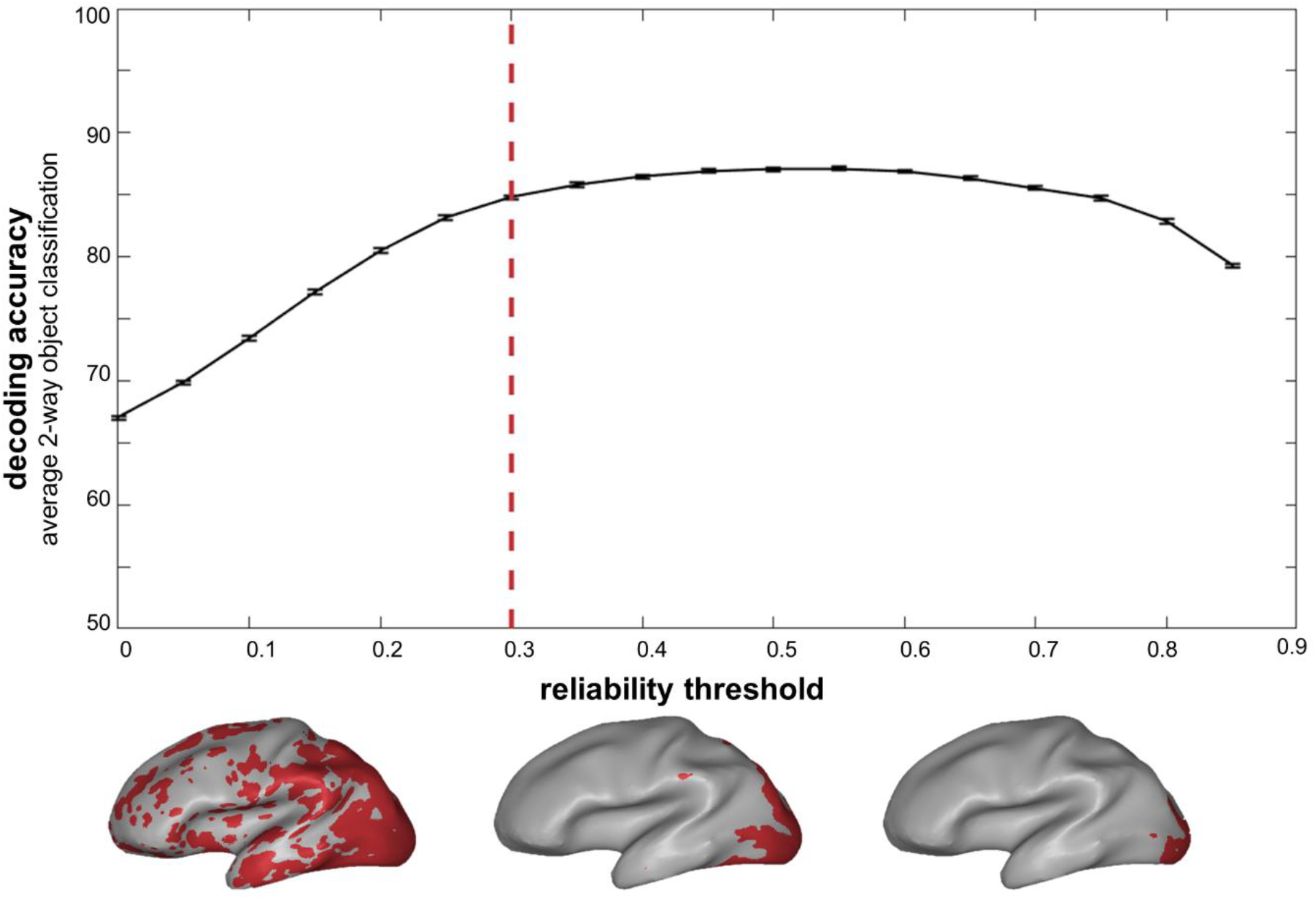
Replication of the Decoding Analysis. Results of pairwise object classification analysis (decoding responses to 72 object images in the replication dataset) are plotted at a range of reliability thresholds. The black line indicates average performance across object pairs, and error bars indicate standard error of the mean. Brain figures show the reliable coverage in the group data at three sample thresholds: *r* = 0.1, 0.4, and 0.7. Red dashed line indicates the final reliability threshold selected for this dataset.

**Figure S2:**
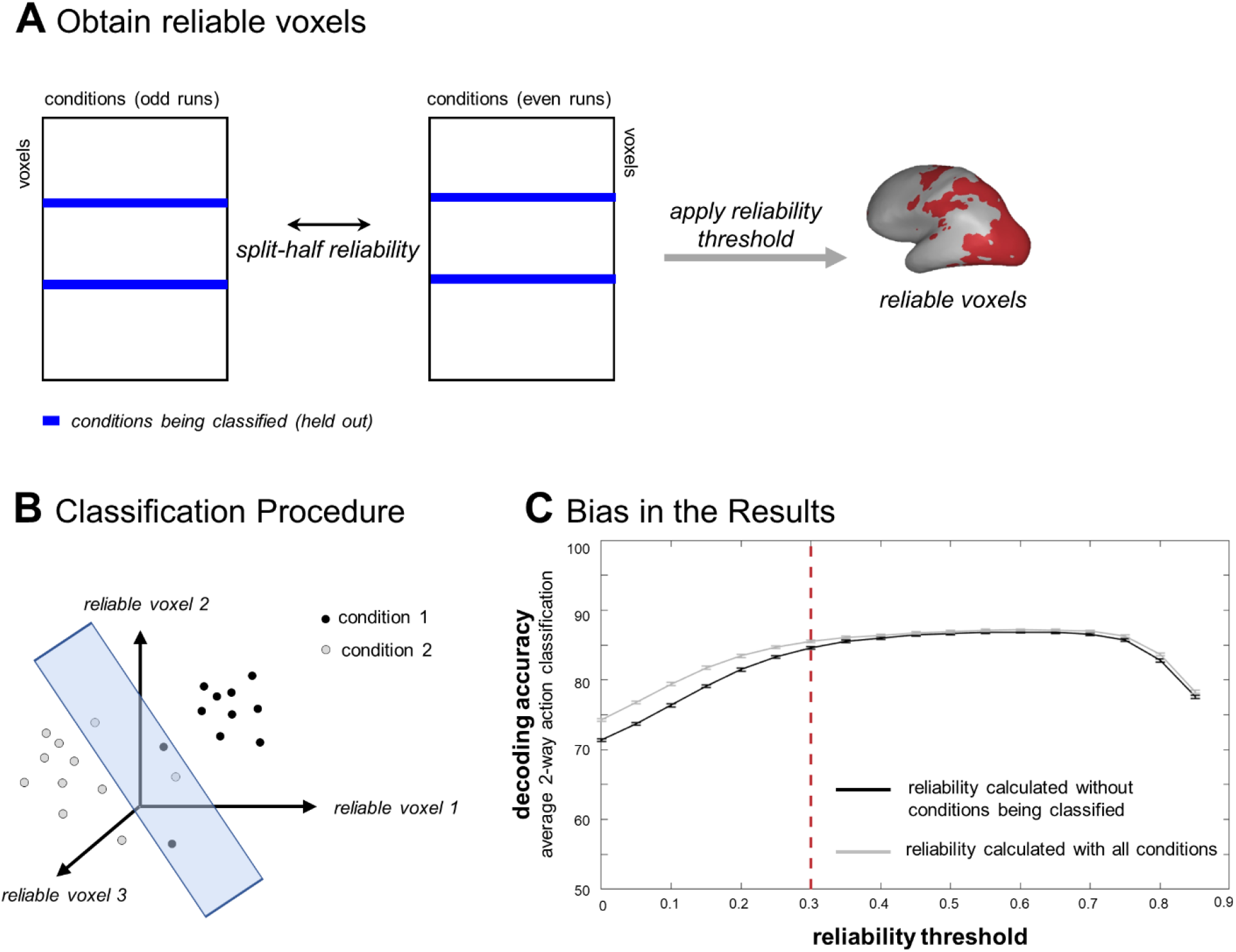
Avoiding Bias in Decoding Analyses. We outline a procedure to perform a pair-wise decoding analysis in reliable voxels without inflating estimates of the decoding accuracy. (A) For each pair of conditions being classified, we defined the reliable voxels <u>*without*</u> the pair that was being classified. As this schematic illustrates, this can be done in two steps: split-half reliability is calculated over the whole brain using the group-level responses to all conditions *except* the two being classified. Then, the reliability threshold is applied to obtain the reliable voxels. Here, we determined this threshold beforehand, using the full dataset; alternatively, it could be calculated uniquely for each loop of this analysis (which would result in a slightly different reliability threshold for each pair of conditions being classified). (B) Once the reliable voxels have been identified, the next step is to train a classifier to distinguish between the two conditions that were held out, based on responses in the reliable voxels. Here, we trained the classifier on 80% of the data, then tested on the remaining 20%. (C) Plot illustrating that defining the reliable voxels using responses to *all* conditions can result in an artificial boost to the classification accuracy, particularly at low reliability thresholds. Dashed red line indicates the threshold used to define reliable voxels in this dataset.

## REFERENCES

Bonnett, D. G., & Wright, T. A. (2000). Sample size requirements for estimating Pearson, Kendall, and Spearman correlations. Psychometrika, 65, 23–28.

Cox, D. D., & Savoy, R. L. (2003). Functional magnetic resonance imaging (fMRI) “brain reading”: detecting and classifying distributed patterns of fMRI activity in human visual cortex. Neuroimage, 19(2), 261–270.

Duncan, J., & Owen, A. M. (2000). Common regions of the human frontal lobe recruited by diverse cognitive demands. Trends in Neurosciences, 23(10), 475–483.

Dunlap, H. F. (1931). An empirical determination of the distribution of means, standard deviations and correlation coefficients drawn from rectangular populations. The Annals of Mathematical Statistics, 2(1), 66–81.

Eklund, A., Nichols, T. E., & Knutsson, H. (2016). Cluster failure: why fMRI inferences for spatial extent have inflated false-positive rates. Proceedings of the National Academy of Sciences, 201602413.

Gayen, A. (1951). The frequency distribution of the product-moment correlation coefficient in random samples of any size drawn from non-normal universes. Biometrika, 38(1/2), 219–247.

Güçlü, U., & van Gerven, M. A. (2015). Deep neural networks reveal a gradient in the complexity of neural representations across the ventral stream. Journal of Neuroscience, 35(27), 10005–10014.

Hanson, S.J., Matsuka, T., Haxby, J.V., 2004. Combinatorial codes in ventral temporal lobe for object recognition: Haxby (2001) revisited: is there a “face” area? Neuroimage 23(1), 156–166.

Hasson, U., Levy, I., Behrmann, M., Hendler, T., & Malach, R. (2002). Eccentricity bias as an organizing principle for human high-order object areas. Neuron, 34(3), 479–490.

Hasson, U., Harel, M., Levy, I., & Malach, R. (2003). Large-scale mirror-symmetry organization of human occipito-temporal object areas. Neuron, 37(6), 1027–1041.

Haxby, J. V., Guntupalli, J. S., Connolly, A. C., Halchenko, Y. O., Conroy, B. R., Gobbini, M. I., Hanke, M., & Ramadge, P. J. (2011). A common, high-dimensional model of the representational space in human ventral temporal cortex. Neuron, 72(2), 404–416.

Haxby, J. V. (2012). Multivariate pattern analysis of fMRI: the early beginnings. Neuroimage, 62(2), 852–855.

Huth, A. G., Nishimoto, S., Vu, A. T., & Gallant, J. L. (2012). A continuous semantic space describes the representation of thousands of object and action categories across the human brain. Neuron, 76(6), 1210–1224.

Huth, A. G., de Heer, W. A., Griffiths, T. L., Theunissen, F. E., & Gallant, J. L. (2016). Natural speech reveals the semantic maps that tile human cerebral cortex. Nature, 532(7600), 453.

Jiang, Y., & Kanwisher, N. (2003). Common neural mechanisms for response selection and perceptual processing. Journal of Cognitive Neuroscience, 15(8), 1095–1110.

Johansen-Berg, H., Behrens, T. E. J., Robson, M. D., Drobnjak, I., Rushworth, M. F. S., Brady, J. M., … & Matthews, P. M. (2004). Changes in connectivity profiles define functionally distinct regions in human medial frontal cortex. Proceedings of the National Academy of Sciences, 101(36), 13335–13340.

Jozwick, K. M., Kriegeskorte, N., & Mur, M. (2016). Visual features as stepping stones toward semantics: Explaining object similarity in IT and perception with non-negative least squares. Neuropsychologia, 83, 201–226.

Kay, K. N., & Yeatman, J. D. (2017). Bottom-up and top-down computations in word- and face-selective cortex. Elife, 6, e22341.

Konkle, T., & Oliva, A. (2012). A real-world size organization of object responses in occipitotemporal cortex. Neuron, 74(6), 1114–1124.

Kriegeskorte, N., Goebel, R., & Bandettini, P. (2006). Information-based functional brain mapping. Proceedings of the National Academy of Sciences, 103(10), 3863–3868.

Kriegeskorte, N., Mur, M., & Bandettini, P. A. (2008). Representational similarity analysis-connecting the branches of systems neuroscience. Frontiers in Systems Neuroscience, 2, 4.

Kriegeskorte, N., Simmons, W. K., Bellgowan, P. S., & Baker, C. I. (2009). Circular analysis in systems neuroscience: the dangers of double dipping. Nature Neuroscience, 12(5), 535.

Lashkari, D., Vul, E., Kanwisher, N., & Golland, P. (2010). Discovering structure in the space of fMRI selectivity profiles. Neuroimage, 50(3), 1085–1098.

Long, B., Yu, C. P., & Konkle, T. (2018). Mid-level visual features underlie the high-level categorical organization of the ventral stream. Proceedings of the National Academy of Sciences, 115(38), E9015–E9024.

Magri, C. Long, B., Chiou, R., & Konkle, T. (May, 2019). Behavioral and Neural Associations between Object Size and Curvature. Poster presented at the annual meeting of the Vision Sciences Society, St. Pete’s Beach, FL.

Mitchell, T. M., Shinkareva, S. V., Carlson, A., Chang, K. M., Malave, V. L., Mason, R. A., & Just, M. A. (2008). Predicting human brain activity associated with the meanings of nouns. Science, 320(5880), 1191–1195.

Mur, M., Meys, M., Bodurka, J., Goebel, R., Bandettini, P. A., & Kriegeskorte, N. (2013). Human object-similarity judgments reflect and transcend the primate-IT object representation. Frontiers in Psychology, 4, 128.

Naselaris, T., Stansbury, D. E., & Gallant, J. L. (2012). Cortical representation of animate and inanimate objects in complex natural scenes. Journal of Physiology-Paris, 106(5-6), 239–249.

Nishimoto, S., Vu, A. T., Naselaris, T., Benjamini, Y., Yu, B., & Gallant, J. L. (2011). Reconstructing visual experiences from brain activity evoked by natural movies. Current Biology, 21(19), 1641–1646.

Norman, K.A., Polyn, S.M., Detre, G.J., Haxby, J.V., 2006. Beyond mind-reading: multivoxel pattern analysis of fMRI data. Trends in Cognitive Science, Sep. 10 (9), 424–430.

Norman-Haignere, S., Kanwisher, N. G., & McDermott, J. H. (2015). Distinct cortical pathways for music and speech revealed by hypothesis-free voxel decomposition. Neuron, 88(6), 1281–1296.

Orlov, T., Makin, T. R., & Zohary, E. (2010). Topographic representation of the human body in the occipitotemporal cortex. Neuron, 68(3), 586–600.

Pearson, E. S. (1931). The analysis of variance in cases of non-normal variation. Biometrika, 23(1/2), 114–133.

Pereira, F., Mitchell, T., & Botvinick, M. (2009). Machine learning classifiers and fMRI: a tutorial overview. Neuroimage, 45(1), S199–S209.

Saxe, R., Brett, M., & Kanwisher, N. (2006). Divide and conquer: a defense of functional localizers. Neuroimage, 30(4), 1088–1096.

Tarhan, L., & Konkle, T. (2019). Sociality and Interaction Envelope Organize Visual Action Representations. bioRxiv, 618272.

Thornton, M. A., & Mitchell, J. P. (2017). Theories of person perception predict patterns of neural activity during mentalizing. Cerebral cortex, 1–16.

